# Live-Cell Imaging and Quantification of PolyQ Aggregates by Stimulated Raman Scattering of Selective Deuterium Labeling

**DOI:** 10.1101/820217

**Authors:** Kun Miao, Lu Wei

## Abstract

Huntington’s disease, a major neurodegenerative disorder, involves deposition of aggregation-prone proteins with long polyglutamine (polyQ) expansions. The ability to non-perturbatively visualize the formation of aggregates could offer new molecular insight for their pathologic roles. Here, we propose stimulated Raman scattering imaging of deuterium-labeled glutamine to investigate native polyQ aggregates in live cells with subcellular resolution. Through the enrichment of deuterated glutamine in the polyQ sequence of mutant Huntingtin (mHtt) proteins, we first achieved sensitive and specific SRS imaging of carbon-deuterium bonds (C-D) from aggregates without GFP labeling. These aggregates become 1.8-fold denser compared to those with GFP. Second, we performed ratiometric quantification, which revealed a dependence of protein compositions on aggregation sizes. Moreover, we calculated the absolute concentrations for sequestered mHtt and non-mHtt proteins within the same aggregates. Our method may readily reveal new features of polyQ aggregates and could be suited for *in vivo* investigations on multicellular organisms.

---

A hallmark of neurodegenerative disorders is the presence of protein aggregates in peripheral nerves^1–4^. Among these disorders are polyglutamine (polyQ) diseases, such as Huntington’s disease (HD), which starts with motor symptoms like chorea and followed by memory deficit and depressions^3,4^. The onset of HD has been linked to abnormally expanded CAG trinucleotide repeats that encode polyQ sequence in mutant Hungtingtin (mHtt) protein. While Q repeats are typically fewer than 37 in healthy humans, they can range from 40 to 250 in Huntington’s patients and are consistently found in the protein depositions of HD brain slices by immunohistology^1,3^. However, the pathological roles of polyQ aggregates still remain elusive^2,4–6^. Recent studies suggested that soluble oligomers are cytotoxic by dynamically interacting with functional proteins and triggering apoptosis while aggregates are cyto-protective by sequestering toxic protein oligomers to form stable inclusion bodies^4,7,8^. In contrast, evidence also indicates that toxicity of aggregates arises from depleting functional (e.g. chaperones and transcription factors) and structural (e.g. actin) proteins and impairing cellular organelles (e.g. ribosomes and endoplasmic reticulum)^9–13^.

To understand their molecular roles, extensive efforts have been made to investigate the compositions, structures, and kinetics of mHtt aggregates. Conventional biochemical assays and recent quantitative proteomics offer relative protein compositions from the aggregates in reference to the soluble pools. However, these methods rely on extensive post-processing such as aggregation purification and solubilization^9,10^. *In vitro* spectroscopic studies including IR^14^, UV-resonance Raman^15,16^, NMR spectroscopy^17^ and fluorescence^18,19^ on model peptides provide crucial information, but they are limited to relatively short expansion lengths because of the difficulty in isolating peptides with long Q repeats^14–19^. More importantly, all these *in vitro* studies cannot recapitulate the native aggregation status in live cells, which have complex intracellular environment. For live-cell studies, fluorescence imaging offers unprecedented spatial and temporal resolution, by fusing fluorescent proteins^20^ or self-labeling tags (e.g. HaloTag)^21^ to the C-terminus of a mHtt exon1 (ex1) sequence (Fig. 1a). The aggregation-prone ex1 fragment, which comprises of a 17 N-terminal sequence, a polyQ tract followed by a proline-rich domain at the C terminus (Fig. 1a), can effectively induce pathological phenotype of HD in transgenic mouse model and humans^4,22^. Compared to mHtt ex1, however, GFP is much larger in size and has a known tendency to oligomerize^23^. This could perturb the aggregation kinetics and conformations, and may contribute to the controversy of reported toxicity. Moreover, because the dense aggregation environment often causes fluorescence quenching^24^, fluorescence imaging doesn’t allow quantitative analysis of aggregates. It is therefore highly desirable to have a new modality that combines the advantages from *in vitro* investigations and fluorescence imaging while overcoming their limitations.

**Fig. 1.**
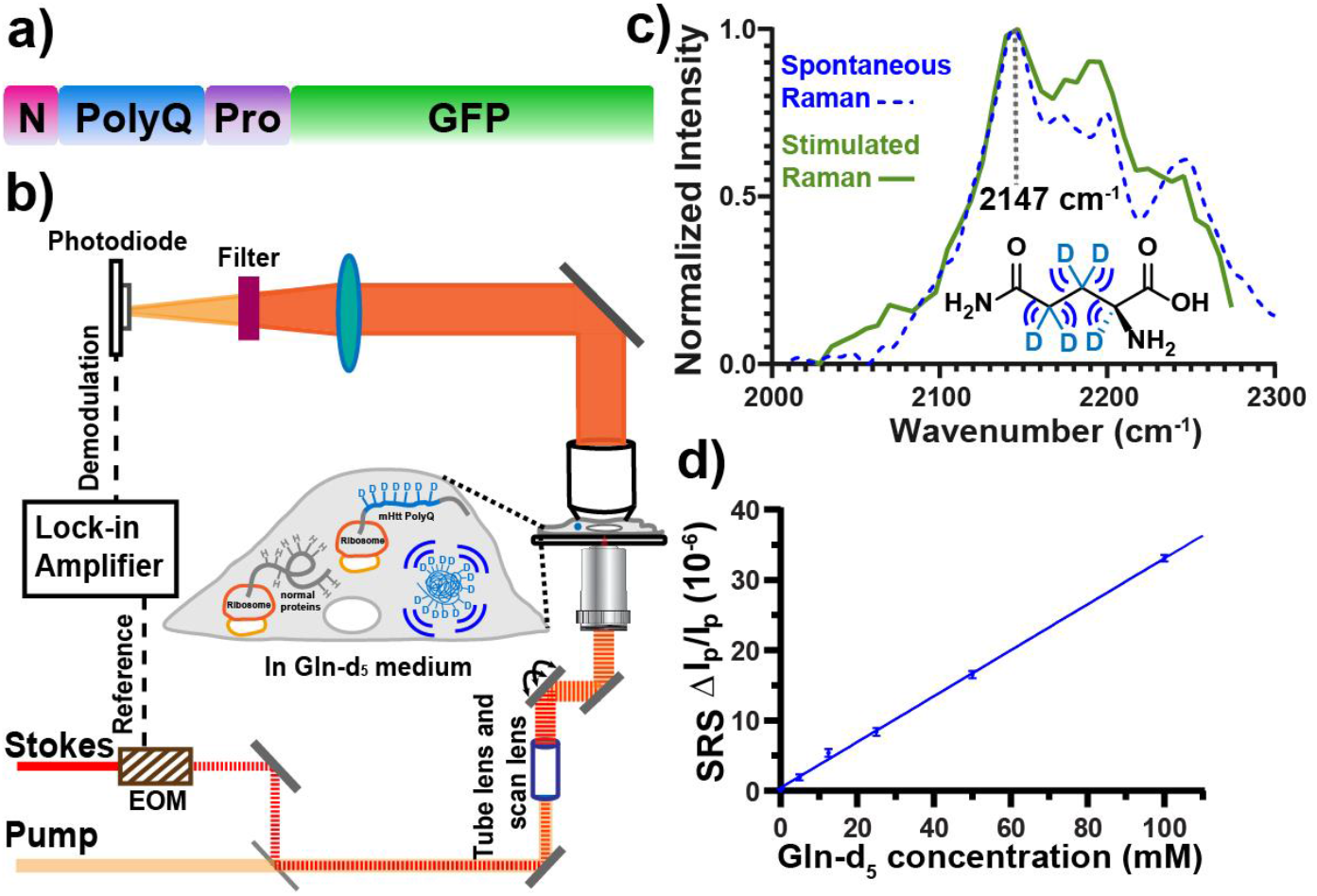
Experimental scheme for stimulated Raman scattering (SRS) microscopy with deuterated glutamine (Gln or Q) labeling. a) Plasmid construct of a model mutant Huntingtin (mHtt) Exon1 (ex1) protein fused with GFP at C terminus. b) Experimental scheme for SRS imaging of Gln-d_5_ labeled polyQ aggregates. c) Spontaneous Raman (blue dashed) and SRS (green) spectra of 60 mM Gln-d_5_ solution. d) Linear dependence of SRS signals (at 2,147 cm^−1^) on Gln-d_5_ concentrations under a 50-μs time constant. Error bar: SD.

Here, we report a general method for live-cell imaging and quantification of polyQ aggregates by stimulated Raman scattering (SRS) microscopy (Fig. S1) of deuterium-labeled glutamine (Gln) (Fig. 1b, c). Labeling of aggregates is achieved through replacing regular Gln in the medium with the Gln-d_5_ (Fig. 1c), which would be metabolically incorporated and enriched into the long polyQ tail of expressed mHtt proteins (Fig. 1a, b). Targeting the vibrational frequency of carbon-deuterium bonds (C-D), SRS imaging obtains subcellular mapping of mHtt aggregates in live cells without GFP, which is typically employed in fluorescence microscopy. In parallel to C-D imaging, SRS allows label-free visualization of endogenous proteins at the CH_3_ vibrational channel (2940 cm^−1^)^25^. While C-D signals represent Gln-d_5_ enriched mHtt proteins, CH_3_ signals mostly originate from non-mHtt cytosolic proteins. This feature enables us to perform ratiometric analysis of CH_3_-to-CD ratios (CH/CDs) to investigate polyQ aggregate compositions. We revealed a decrease of CH/CDs as the aggregation sizes increase. Finally, combining aggregation-specific CH/CDs with calibrated CD intensities, we calculated molar percentages and absolute concentrations for mHtt and non-mHtt proteins from polyQ aggregates in live cells. Such quantification is otherwise highly challenging for existing methods.

The coupling of SRS imaging with Gln-d_5_ labeling presents following advantages. First, the vibrational frequency of C-D in Gln-d_5_ (Fig. 1c) is in the desired cell-silent region (1800-2600 cm^−1^), providing high imaging specificity without background from endogenous biomolecules. Second, the non-exchangeable labeling on the side chains offers reliable signals. Third, Gln enrichment in the polyQ region (Figs. 1a) offers both high labeling specificity and SRS imaging sensitivity. For example, for a widely used mHtt-97Q ex1 fragment, Gln accounts for 68% of the ex1 sequence (Fig. S2), while the natural occurrence of Gln is only about 4.2% in human proteomes^26^. Compared to label-free SRS^25^ and SRS imaging with ^13^C-Phenyalanine^27^ or deuterated all essential amino acids^28^, selective Gln-d_5_ labeling is hence more specific for imaging polyQ aggregates. Fourth, compared to using alkyne-tagged unnatural amino acids, which only introduces one tag to one copy of a protein^29^, multiple Q labeling (e.g. 103Q for mHtt-97Q ex1) offers higher sensitivity and less sample manipulation. Fifth, compared to spontaneous Raman^30^, SRS offers higher detection sensitivity and faster image acquisition with stimulated emission quantum amplification principle (Fig. S1). Compared to Coherent anti-Stokes Raman scattering (CARS)^31^, another nonlinear microscopy technique, SRS provides high-fidelity Raman spectra (Fig. 1c) and linear concentration dependence (Fig. 1d) without non-resonance background.

We first determined our SRS detection limit on Gln-d_5_ solution to be 3 mM (Fig. 1d, when signal (S)/noise (N)=1) by targeting the C-D peak at 2147 cm^−1^. Hence, our detection limit is as low as 29 μM for mHtt proteins with a total of 103 Gln in the mHtt-97Q ex1 protein. This strategy offers higher detection sensitivity compared to previously reported 200 μM by SRS imaging of alkyne tags^32^. In cells, however, the detection limit is set by aggregation-signal to cellular-background ratio (S_Agg_/B_Cell_). There are two sources for B_Cell_. First, the newly synthesized proteome incorporates Gln, which only accounts for 4.2 % of human proteome^25^. The estimated detectability for mHtt-97Q ex1 is 86 μM when S_Agg_/B_Cell_=1 (details in SI). Second, the intracellular free Gln pool is about 8 mM^33^, slightly lowering the achievable detectability. To minimize this additional background, we replaced the medium with buffer before imaging.

We next validated our SRS imaging by transfecting HeLa cells with mHtt-97Q-GFP plasmid (Figs. 1a and S2) and culturing them in Gln-d_5_ medium. We first conducted parallel SRS and fluorescence imaging on the same set of live cells. The SRS image of C-D enriched aggregate (Fig. 2a, 2167 cm^−1^) agrees well with the fluorescence image of GFP (Fig. 2a, fluorescence). A clear off-resonance image demonstrates high SRS imaging quality (Fig. 2a, 2035 cm^−1^). The C-D peak for Gln-d_5_ is shifted from 2147 cm^−1^ in solution to 2167 cm^−1^ after being incorporated into cellular proteins, suggesting a change of microenvironment. We found that the shifted C-D spectrum has Raman spectral features from both Gln-d_5_ solution and solid (peaked at 2167 cm^−1^) (Fig. S3). Label-free SRS images at CH_3_ (2940 cm^−1^) and Amide I (1664 cm^−1^) channels, widely adopted for imaging total proteins^27^, show much decreased detection specificity. In particular, these decreases were observed for small-size aggregates (Fig 2b, arrow indicated), which are indistinguishable from the nucleoli. Moreover, the high-contrast of C-D image decreases significantly if Gln-d_5_ is replaced by non-enriched Leucine-d_10_ (Leu-d_10_) for labeling (Fig. 2c). Our quantification on the average S_Agg_/B_Cell_ for each channel clearly demonstrated much higher imaging specificity with Gln-d_5_ labeling (Fig. 2d).

**Fig. 2.**
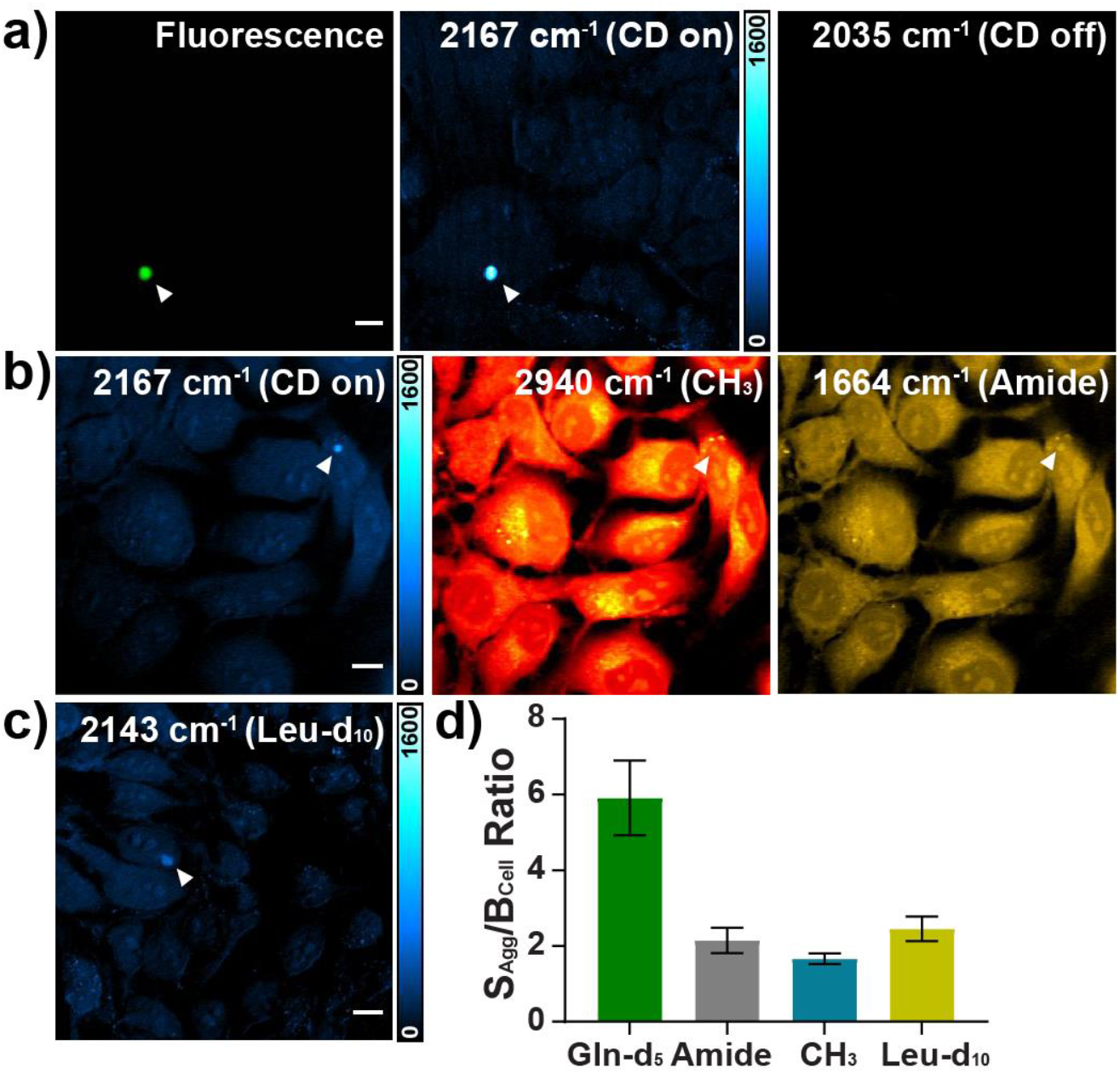
Live-cell SRS imaging of mHtt-97Q-GFP aggregates with Gln-d_5_ labeling. a) SRS imaging of mHtt aggregates (arrowheaded, 2167 cm^−1^, C-D on), validated by fluorescence imaging through GFP (Fluorescence). Off-resonance image at 2035 cm^−1^ shows no signal. b) Live-cell SRS images for an mHtt-97Q-GFP aggregate (arrowheaded) at Gln-d_5_ (2167 cm^−1^), CH_3_ (2940 cm^−1^) and Amide I (1664 cm^−1^) channels on the same set of Hela cells. c) SRS imaging of an mHtt-97Q-GFP aggregate at 2143 cm^−1^ by Leucine-d_10_ (Leu-d_10_) labeling. d) Average S_Agg_/B_cell_ from SRS images of C-D with Gln-d_5_ labeling (5.75±1.03, n=13); Amide I (2.15±0.34, n=4); CH_3_ (1.66±0.14, n=10) and C-D with Leu-d_10_ labeling (2.45±0.33, n=10.) Error bar: SD.

After establishing the feasibility of SRS imaging of Gln-d_5_ labeled mHtt-97Q-GFP aggregates, we aim to image native mHtt-97Q proteins without GFP (Fig. 3a). GFP may perturb aggregation formation of mHtt proteins, because GFP is 238 amino acid (aa) in length, which is about twice as large as mHtt-97Q ex1 with only 152 aa. We successfully imaged aggregates at the same C-D frequency (Fig. 3b, CD on and off). Interestingly, these aggregates (Fig. 3b, 2167 cm^−1^) are significantly brighter than those with GFP labeling (Fig. 2a, 2167 cm^−1^). Because GFP sequence only contain 8 Gln, the C-D intensity should remain approximately unchanged when deleting GFP. We hence reasoned that the detected intensity increase is due to the formation of denser aggregates. A similar phenomenon was recently reported by Cryo-ET with the same mHtt ex1 sequence^11^. To test our theory, we acquired Amide I images to compare total protein concentrations between aggregates with and without GFP. If the density remains unchanged, mHtt-97Q aggregates would have a much lower amide intensity than that of mHtt-97Q-GFP because the GFP sequence contributes significantly to amide signals. We observed similar levels of amide signals between these two types of aggregates (Fig. S4), indicating that the increase in aggregate density makes up for the loss of GFP. Moreover, mHtt-97Q aggregates become barely distinguishable from cellular background in CH_3_ channel (Fig. 3b, 2940 cm^−1^), suggesting that previous CH_3_ signals for aggregates were largely from GFP sequence (Fig. 2b, 2940 cm^−1^ and Fig. S2). To quantify aggregates from multiple experimental replicates, we plotted the average S_Agg_/B_Cell_ in both C-D and CH channels for mHtt-97Q and mHtt-97Q-GFP aggregates (Fig. 3c). In C-D channel, the S_Agg_/B_Cell_ for mHtt-97Q aggregates is 1.8-fold higher, confirming the formation of 1.8-fold denser aggregates without GFP labeling. In C-H channel, the S_Agg_/B_Cell_ values remain approximately the same.

**Fig. 3.**
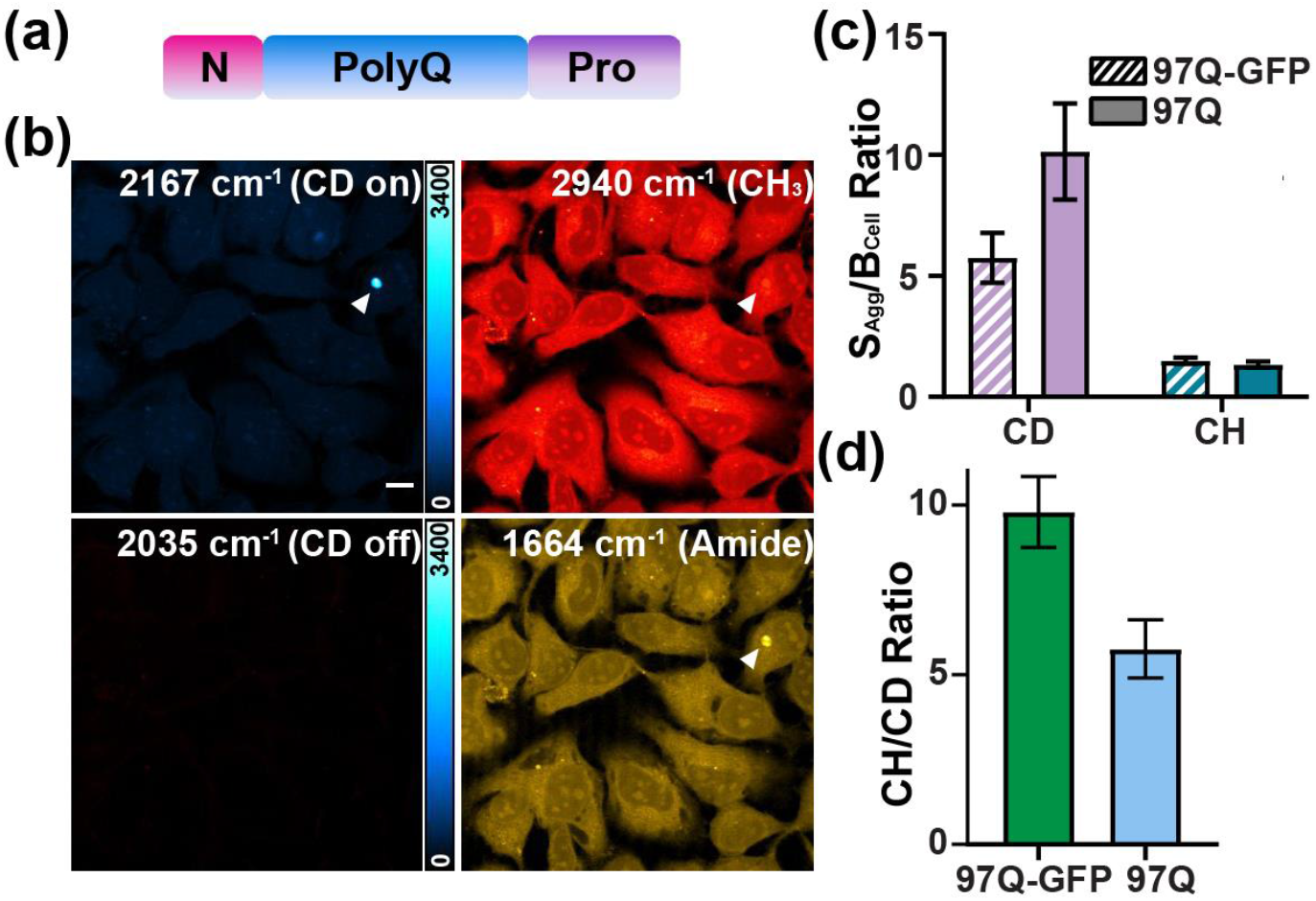
SRS imaging of mHtt-97Q aggregates without GFP. a) Plasmid construct of mHtt-97Q by deleting GFP sequence in Fig. 1a. b) SRS images of a Gln-d_5_ labeled aggregate at C-D on-resonance (2167 cm^−1^), C-D off-resonance (2035 cm^−1^), CH_3_ (2940 cm^−1^) and Amide I (1664 cm^−1^). c) S_Agg_/B_Cell_ of mHtt-97Q (97Q) in C-D (CD, 10.14±1.99, n=11) and CH_3_ (CH, 1.50±0.12, n=10) channels compared to that for mHtt-97Q-GFP (97Q-GFP, CD: 5.75±1.03, n=13; CH: 1.66±0.14, n=10). d) CH/CDs for 97Q-GFP (9.79±1.05, n=7) and 97Q (5.75±0.86, n=20) aggregates. Scale bar: 10 μm. Error bar: SD.

With multi-channel SRS imaging, we next performed live-cell ratiometric quantification of CH/CDs on the same aggregates. The CH_3_ signals at 2940 cm^−1^ represent non-Gln aa, which mostly come from the sequestered non-mHtt proteins, and the CD signals are from Gln-d_5_, which largely originates from mHtt proteins. The CH/CDs could hence be used as an indicator for aggregation compositions of non-mHtt and mHtt proteins. We first adopted CH/CD quantification to mHtt-97Q and mHtt-97Q-GFP aggregates (Fig. 3d). Surprisingly, the average CH/CD is 5.75 for mHtt-97Q aggregates (Fig. 3d). If aggregates were formed primarily by mHtt proteins, the CH/CD should be about 0.5 because mHtt-97Q sequence contains 49 non-Q aa and 103 Q. Our control experiments confirmed that Gln-d_5_ labeling efficiency is close to 100% (i.e. *de novo* glutamine synthesis for regular Gln is about zero, Fig. S5 and S6), indicating all 103 Q are deuterated. Similarly, the CH/CD should have decreased by 5 folds from mHtt-97Q-GFP (289 non-Q aa and 112 Q) to mHtt-97Q aggregates (49 non-Q aa and 103 Q) (Fig. S2). Instead, the decrease is only 1.5-fold (Fig. 3d, 9.79 vs 5.75). Our results imply that these aggregates contain a rather high percentage of non-mHtt proteins. Sequestrations and depletions of cellular functional and structural proteins by aggregates have been suggested to be one underlying mechanism of HD cytotoxicity^9–13^. We next aim to calculate the relative molar percentages for mHtt and non-mHtt proteins in the aggregates, assuming that the molar percentage of mHtt proteins in the aggregates is x, and then non-mHtt proteins is 1-x. With CH/CDs (Fig. 3d), we can generate an equation with x and (1-x) for the relative ratio between non-Gln aa and Gln-d_5_ (details in SI). We calculated that the average molar percentage of non-mHtt proteins is about 50% (details in SI), which is indeed significant.

Now that we have established CH/CDs could serve as a direct indicator of aggregation compositions, we next examined the relationship between CH/CDs and aggregation sizes. Based on the sequestration theory^12,13^, we expect that the relationship might offer insight into understanding the molecular cytotoxicity of polyQ aggregates. Interestingly, we observed a negative correlation (Fig. 4a, Pearson’s r=−0.56), which indicates that the percentage of sequestered mHtt proteins increases as the aggregates grow. As a comparison, such distinctive correlation by Gln-d_5_ labeling is absent when replaced by Leu-d_10_ labeling (Fig. 4b, Pearson’s r=−0.11), because mHtt and non-mHtt proteins have similar Leucine abundance. This underscores the importance of specific polyQ labeling to observe such a size-composition effect.

**Fig. 4.**
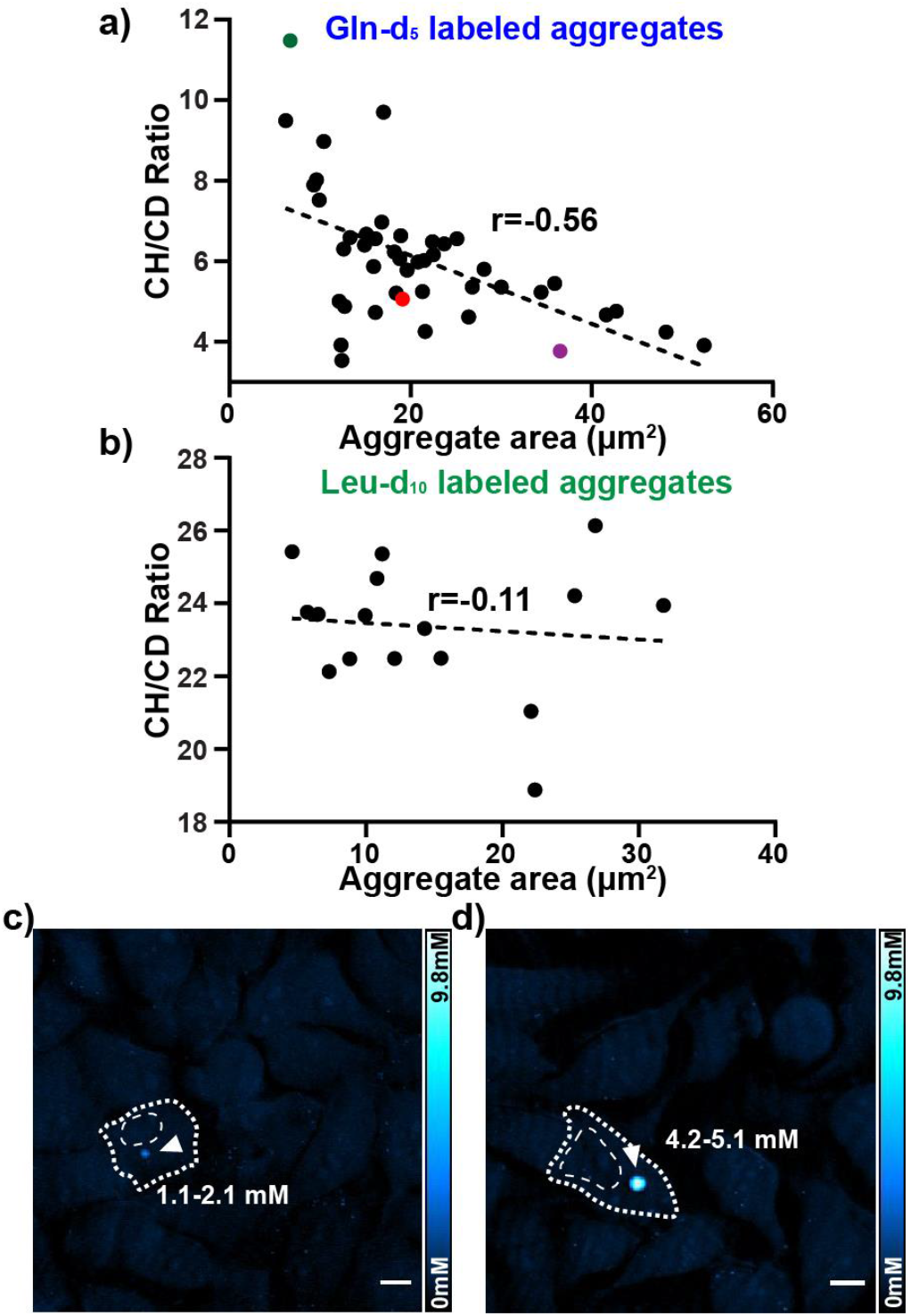
SRS quantification for mHtt-97Q aggregates of different sizes. a) CH/CDs for Gln-d_5_ labeled aggregates present a negative correlation on aggregation areas (The Pearson’s coefficient r=−0.56). b) Minimum correlation between CH/CDs for Leu-d_10_ labeled aggregates and aggregation areas (The Pearson’s coefficient r=−0.11). c, d) C-D SRS images for representative Gln-d_5_ labeled small (c, 6.7 μm^2^, green dot in a) and medium (d, 19.1 μm^2^, red dot in a) mHtt-97Q aggregates. Calculated mHtt-97Q concentrations for the aggregates are indicated. Cell shapes and nuclei are outlined by white dotted line. Scale bar: 10 μm.

In addition to calculating molar percentages from CH/CDs, we can also compute absolute concentrations with aggregate intensities in C-D channel and our reference calibration curves (Fig. 1d). We selected three representative aggregates with different sizes for calculation (Fig. 4a, color-indicated, details in SI and results listed in Table S1). The mHtt concentration in the small aggregate (Fig. 4a, green dot) ranges from 1.2 - 2.1 mM (Fig. 4c, 6.7 μm^2^). It increases to 4.3 - 5.1 mM (and 4d, 19.1 μm^2^) for the medium aggregate (Fig. 4a red dot). The concentration becomes 5.6 - 6.4 mM for the large aggregate (36.5 μm^2^, Fig. 4a, magenta dot). The upper and lower limits shown are determined by the relative content of newly synthesized and pre-existing non-mHtt proteins sequestered by the aggregates. Surprisingly, we found that while mHtt concentrations increase with sizes, the concentrations of non-mHtt proteins remain almost the same for small (4.6 mM), medium (4.7 mM) and large (4.2 mM) aggregates. Our observations suggest that the formations of small aggregates preferentially sequester non-mHtt cytosolic proteins. These proteins are likely functional chaperones, ribonucleoproteins and structural proteins.^9–13^ As the aggregates become larger, they then sequester more mHtt proteins. This might indicate a cellular rescue mechanism by clearing toxic mHtt proteins^4,7,8^. Our data further demonstrate that the total protein concentrations of these aggregates fall in the range of 5-10 mM. To the best of our knowledge, our study is the first to quantify the mHtt protein concentrations and molar percentages for aggregates of different sizes in live cells.

In summary, we combined SRS microscopy with Gln-d_5_ labeling to achieve sensitive and specific imaging as well as ratiometric quantification of polyQ aggregations in HD. Our technique is applicable to polyQ expansions of various lengths. In particular, with linear concentration dependence, it is suited for investigating aggregates of extended polyQ construct (e.g. >200Q), which may form rather dense structures and pose challenges to study by other strategies. Our method is also applicable to other polyQ diseases, including spinocerebellar ataxia and spinobulbar muscular atrophy^34^. Other poly-aa diseases^35,36^ with poly-GA, , poly-PR, and poly-PA aggregates, recently reported in ALS/FTD patient brain, could also be investigated by selective deuteration of corresponding aa. In addition, since Gln can transport across blood-brain barrier^37^ and deuterium labeling is minimally invasive, applications to animal models or even to humans may be possible. Moreover, correlative live-cell SRS imaging with Gln-d_5_ labeling with recently demonstrated Cryo-ET^11^ or quantitative proteomics^12^ may offer comprehensive structure-function relationship for native mHtt-97Q aggregates.

## Acknowledgments

We would like to thank Dr. W. Min, Dr. C. Qian, D. Lee, J. Du and L. Voong for helpful discussion. We are grateful for the plasmid (mHtt-97Q-GFP) shared by Prof. R. Kopito and Prof. F.-U. Hartl. L. Wei acknowledges the support of start-up funds from California Institute of Technology.

