## Supplementary Information for "Live-Cell Imaging and Quantification of PolyQ Aggregates by Stimulated Raman Scattering of Selective Deuterium Labeling"

### for

#### Experimental Methods

**Stimulated Raman Scattering (SRS) Microscopy** A picoEmerald laser system (Applied Physics and Electronics) was used as the light source for SRS microscopy. It produces 2 ps pump (tunable from 770 nm – 990 nm, bandwidth 0.5 nm, spectral bandwidth ~ 7 $cm^{-1}$) and Stokes (1031.2 nm, spectral bandwidth 10 $cm^{-1}$) beams with 80 MHz repetition rate. Stokes beam is modulated at 20 MHz by an internal electro-optic modulator. The spatially and temporally overlapped Pump and Stokes beams are introduced into an inverted laser-scanning microscope (FV3000, Olympus), and then focused onto the sample by a 25X water objective (XLPLN25XWMP, 1.05 N.A., Olympus). Transmitted Pump and Stokes beams are collected by a high N.A. condenser lens (oil immersion, 1.4 N.A., Olympus) and pass through a bandpass filter (893/209 BrightLine, 25mm, Semrock) to filter out Stokes beam. A large area (10×10 mm) Si photodiode (S3590-09, Hamamatsu) is used to measure the pump beam intensity. A 64 V reverse-biased DC voltage is applied on the photodiode to increase the saturation threshold and reduce response time. The output current is terminated by a 50Ω terminator and pre-filtered by a 19.2-23.6-MHz band-pass filter (BBP-21.4+, Mini-Circuits) to reduce laser and scanning noise. The signal is then demodulated by a lock-in amplifier (SR844, Stanford Research Systems) at the modulation frequency. The in-phase X output is fed back to the Olympus IO interface box (FV30-ANALOG) of the microscope. 30 µs time constant is set for the lock-in amplifier. Correspondingly, 80 µs pixel dwell time is used, which gives a speed of 8.5 s/frame for a 320-by-320-pixel image. For 1664, 2035, 2167 and 2940 $cm^{-1}$, the wavelengths of pump laser are 880.6, 852.3, 842.8, and 791.3 nm, respectively. On-sample pump and modulated Stokes beam powers are 30 mW and 45 mW respectively for 1664 and 2940 $cm^{-1}$; 30 mW and 170 mW respectively for 2035, 2143, 2167 $cm^{-1}$. Laser powers are monitored throughout image acquisition by an internal power meter and power fluctuations are controlled within 1%. 16-bit greyscale images are acquired by Olympus Fluoview 3000 software. All data presented are from at least three independent experiments.

**Spontaneous Raman Spectroscopy** Spontaneous Raman spectra were acquired using an upright confocal Raman spectrometer (Horiba Raman microscope; Xplora plus). A 532 nm YAG laser is used to illuminate the sample with a power of 12 mW on sample through a 100×, N.A. 0.9 objective (MPLAN N; Olympus). Data acquisition was performed with 80 s integration by the LabSpec6 software. For Glutamine solution, background was subtracted by measuring signal from non-solution region on the same sample. The spectra are normalized to 2147 $cm^{-1}$.

Same Raman cross-sections for C-D and C-H are confirmed by Spontaneous Raman measurements on pure DMSO and DMSO-$d_6$. Same system throughput of pump lasers at two designated wavelengths for C-H and C-D SRS imaging are confirmed by SRS acquisitions on pure DMSO and DMSO-$d_6$ at the C-H and C-D channels.

**Plasmid construction and molecular cloning.** The plasmid pcDNA3.1-N-httQ97-GFP was a generous gift from Prof. Ron Kopito and Prof. F.-U. Hartl. L. Hartl. The Htt97Q was obtained by deleting the GFP sequence from the pcDNA3.1-N-httQ97-GFP plasmid. The subcloning was done by GenScript USA Inc. For plasmid amplification, mHtt97Q-GFP and mHtt97Q were transformed into DH5α *Escherichia coli* cells. The cells were plated onto agar plates with respective antibiotics for selection for 14h. The resulting single colony was picked and grown in LB medium for 24 h. The cultures were collected, and the plasmids were purified by QIAGEN Maxi-prep kit.

**Cell culture, transfection, metabolic labeling and imaging** Deuterated glutamine DMEM was made by supplying glutamine-$d_5$ (Cambridge Isotope) to glutamine deficient DMEM (Gibco). Deuterated leucine DMEM was made by supplying leucine-$d_{10}$ (Cambridge Isotope) and regular methionine (Sigma-Aldrich) to leucine and methionine deficient DMEM (Thermo Scientific). Deuterated glucose DMEM was made by supplying $d_7$-glucose (Cambridge Isotope) to glucose deficient DMEM (Gibco). The solutions were sterile filtered by 0.22 µm low protein binding filter system (Corning). The filtered solutions were added with 10% FBS and 1% penicillin-streptomycin (Sigma-Aldrich) to make the complete media. Cultured HeLa-CCL2 (ATCC) cells were seeded onto 14 mm glass-bottom microwell dishes (MatTek Corporation) or coverslips (12mm, #1.5, Fisher) for 24 h prior to transfection. Cells were first grown in regular DMEM complete medium until they reached 70-90% confluence. The medium was switched to the designated deuterated medium immediately before transfection. Transfection of 1-µg plasmids encoding mHtt-97Q-GFP or mHtt97Q was performed using Lipofectamine 3000 transfection reagent (Thermo Fisher). After 24h of protein expression, medium was switched to the pre-warmed DPBS buffer before imaging. Coverslips were collected and attached to a microscope slides (1mm thick, VWR) with imaging spacer (Sigma-Aldrich); and the glass-bottom dishes were covered with another coverslip (22x22 mm, #1.5, VWR) on the top. Confocal fluorescence images were obtained by the Olympus FluoView™ FV3000 confocal microscope with SRS setup described above.

**Treatment of Glutamine Synthetase Inhibitor in HeLa cells.** Glutamine synthetase inhibitor L-Methionine sulfoximine (MSO) (CAS#15985-39-4) was purchased as solid from ACROS Organic. The white solid was dissolved in $ddH_2O$ to obtain 300 mM stock solution. Right before transfection, the stock solution was diluted 100 times with Gln-$d_5$ DMEM to reach final concentration of 3mM in DMEM[1]. The HeLa cells seeded onto coverslip or imaging dish were switched to MSO medium immediately before transfection and was incubated with the presence of MSO throughout the 24 h expression time. Before imaging, the MSO medium was changed to pre-warmed DPBS buffer.

**Image processing and data analysis** Color-coding and intensity profile for all images were done by ImageJ. Spectral plotting and signal to background ratio calculations were performed in Graphpad Prism 8.1.1. Figures were assembled in Abode Illustrator.

**Estimation of the SRS detection limit for Gln-$d_5$ labeled mHtt-97Q ex1 proteins**

1) Signal-to-Noise ratio (S/N) limited:

From our concentration curve (Fig. 1d), our detection limit for Gln-$d_5$ solution c(Gln-$d_5$) is 3 mM when S/N=1. There is a total of 103 glutamine in the mHtt97Q exon1 sequence. The detection limit for mHtt-97Q protein $c(mHtt)_{S/N}$ is:

$$c(mHtt)_{S/N} = \frac{c(Gln - d_5)}{\mathrm{n(Q)}} = \frac{3\ mM}{103} = 29\ \mu M$$

2) Aggregation-signal to cellular-background ratio ($S_{Agg}/B_{Cell}$) limited:

The background signal originates from the labeling of glutamine residues in newly synthesized cytosolic proteins by Gln-$d_5$. The average concentration of cytosolic proteins is 2 mM[2,3] and the average length of proteins in eukaryotes is 438 amino acids[4]. Glutamine makes up 4.2 percent[4] of the overall proteome and the cellular proteome synthesis rate is about 1% per hour[5]. In our case, we incubated the cell in Gln-$d_5$ medium for 24h for labeling. The detection limit is hence set by $S_{Agg}/B_{Cell}$=1 when the concentration of Gln-$d_5$ in mHtt proteins (n(Q)=103) is equal that in cytosolic proteins. The mHtt-97Q protein concentration ($c(mHtt)_{Agg/Cell}$) is calculated by:

$$c(mHtt)_{S_{Agg}/B_{cell}} = \frac{c(cytosolic\ Gln{-}d_5)}{n(Q)} = \frac{2mM{*}438aa\ per\ protein{*}1\%{*}24h{*}4.2\%}{103\ Q\ per\ Htt} = \frac{8.83\ mM}{103} = 86\mu M$$

**Compositional analysis of mHtt aggregates (Figs. 3d and 4a-b).**

With measured CH/CD ratios through Gln-$d_5$ labeled aggregates, we can calculate the molar percentages of sequestered mHtt and non-mHtt proteins in the aggregates. Since we supply Gln-$d_5$ in DMEM for metabolic incorporation, the Gln-$d_5$ would unbiasedly label newly synthesized mHtt proteins and endogenous non mHtt proteins. The labeled (newly synthesized) non-mHtt proteins will contribute to C-D signals. We here discuss two extreme scenarios, in which the sequestered non-mHtt cytosolic proteins are either all newly synthesized proteins or pre-existing proteins.

<u>1) If the sequestered cytosolic non-mHtt proteins are all newly synthesized proteins:</u>

Because the newly synthesized cytosolic proteins are labeled by Gln-$d_5$, they could contribute to the measured C-D channel SRS signals from the aggregates. First, on average, there are 6 C-H bonds on a side-chain of each amino acid. This is calculated by summing over the products of the number of side-chain C-H bonds from each amino acid and the relative abundance of this amino acid in human proteome[4]. Second, the average cytosolic proteins contain about 438 amino acids (aa)[4]. Third, Glutamine accounts for 4.2% in human proteome[4]. Fourth, the CH Raman cross-section is the same to CD[6,7] (confirmed by DMSO-$d_6$ and DMSO). Fifth, *de novo* glutamine synthesis is negligible in our experiments (see below Figs. S5 & S6),

Assuming there are $x$ amount (molar percentage) of mHtt and (1- $x$) amount of other cytosolic non-mHtt proteins in the aggregates, we could have the following equation based on our measured CH/CD ratio:

$$\frac{CH}{CD} = \frac{438*(1-x)*(1-4.2\%)+49x}{103x+438*(1-x)*4.2\%} * \frac{6}{5} = 5.75 \quad (1)$$

Here, CH signal comes from CH bonds of non-Gln aa of both non-mHtt proteins and mHtt proteins. CD signal comes from C-D bonds in both Gln-$d_5$ labeled mHtt proteins and non-mHtt proteins. As stated above, average cytosolic proteins contain 438 aa, in which 4.2% percent are Gln and (1-4.2%) are non-Gln aa. In contrast, mHtt proteins have 103 Gln (Q) and 49 non-Gln aa. The normalized concentration of Non-Gln aa in the aggregates is hence: 438*(1-x)*(1-4.2%)+49*x. Similarly, the normalized concentration of Gln-$d_5$ in the aggregates is: 103x+438*(1-x)*4.2%. Since non-Gln aa have an average of 6 side chain C-H and Gln-$d_5$ has 5 labeled side chain C-D, a factor of 6/5 is multiplied above. From Eq.(1), the molar percentage of mHtt proteins in aggregates, x, was solved to be **42.7%**, and the molar percentage of sequestered non mHtt proteins in aggregates is **57.3%**.

2) If the sequestered cytosolic non-mHtt proteins are all pre-existing proteins:

The sequestered non-mHtt proteins are not labeled by Gln-$d_5$ and contribute to no C-D signal. The C-D SRS signals only come from Gln-d5 labeling in the mHtt proteins. All Gln in sequestered non-mHtt proteins only contain C-H side chains. Hence, the CH/CD ratio should be:

$$\frac{CH}{CD} = \frac{438*(1-x)+49x}{103x} * \frac{6}{5} = 5.75 \quad (2)$$

From Eq.2, the molar percentage of mHtt protein in aggregates, x, was solved to be **49.6%**, and the molar percentage of sequestered non-mHtt protein in aggregates is **50.4%**.

Described above are two extreme conditions for sequestered cytosolic non-mHtt proteins. As the sequestered proteins should contain both pre-existing proteins and newly synthesized proteins, the real composition of the mHtt aggregation can be calculated from:

$$\frac{CH}{CD} = [A * \frac{438*(1-x)+49x}{103x} + B * \frac{438*(1-x)*95.8\%+49x}{103x+438*(1-x)*4.2\%}] * \frac{6}{5} = 5.75 \quad (3)$$

where A+B=1. Hence, the two extreme conditions discussed above provide the lower and the upper limits of the molar percentage of mHtt proteins in aggregates, ranging from 42.7% to 49.6%. Respectively, the molar percentage of sequestered non-mHtt proteins ranges from 50.4% to 57.3%. On average, the molar percentage of mHtt proteins is **46%** and that of non-mHtt proteins is **54%**.

**Absolute concentrations of mHtt and non-mHtt proteins in the aggregates (Fig. 4 c,d).**

To calculate absolute concentrations, we define two variables: 1) $x$ for the molar percentage of mHtt proteins in the aggregates, as defined above; and 2) $y$ for the absolute concentration of total proteins in the aggregates. The absolute concentrations of mHtt and non-mHtt proteins are then x*y and (1-x)*y respectively.

We take an aggregate, indicated as the green point in Fig. 4a and shown in Fig 4c, as one example for our calculation. We then selected three representative aggregates of different sizes from Fig. 4a and listed all calculated results in Table S1 below.

Based on the C-D signals of aggregates and the concentration curve (Fig. 1d), generated under the same pump and Stokes powers, we measured the average C-D SRS intensity (SRS (C-D)) of the aggregate to be 478.1 a.u., which corresponds to a Gln-$d_5$ concentrating of 212.2 mM. The CH/CD ratio for this aggregate is 11.5. Similarly, we calculated the upper and the lower concentration limit based on the two extreme conditions.

1) If the sequestered cytosolic non-mHtt proteins are all newly synthesized proteins labeled by Gln-$d_5$, then we have:

$$\frac{CH}{CD} = \frac{438*(1-x)*(1-4.2\%)+49x}{103x+438*(1-x)*4.2\%} * \frac{6}{5} = 11.5 \quad (4)$$

$$x * y * 103 + (1-x) * y * 438 * 4.2\% = 212.2\, mM \quad (5)$$

Solving Eq. (4) and (5), we got x=20.6% and y=5.9 mM. As a result, the concentrations (and the molar percentages) of mHtt and non-mHtt proteins in this small aggregate are 1.2mM (20.6%) and 4.7mM (79.4%) respectively.

2) If the sequestered non-mHtt proteins are all pre-existing proteins without Gln-$d_5$ labeling, we have:

$$\frac{CH}{CD} = \frac{438*(1-x)+49x}{103x} * \frac{6}{5} = 11.5 \quad (6)$$

$$y * 103x = 212.2\, mM \quad (7)$$

Solving Eq. (6) and (7), we got x=31.8% and y=6.5 mM. The concentrations (and the molar percentages) of mHtt and non-mHtt proteins in this small aggregate are 2.1mM (31.8%) and 4.4mM (68.2%) respectively.

As we indicated above, these numbers set the upper and lower limits. The concentration of mHtt proteins in this small aggregate ranges from 1.2 mM to 2.1 mM and the concentration of non-mHtt proteins ranges from 4.4 to 4.7 mM.

| The Small Agg Green dot in Fig. 4a | If the sequestered non-mHtt proteins are all newly synthesized proteins | If the sequestered non mHtt proteins are all pre-existing proteins |
|---|---|---|
| Molar percentage of mHtt | 20.6% | 31.8% |
| Absolute mHtt conc. | 1.2 mM | 2.1 mM |
| Molar percentage of other proteins | 79.4% | 68.2% |
| Sequestered non-mHtt protein conc. | 4.7 mM | 4.4 mM |
| **The medium Agg Red dot in Fig. 4a** | | |
| Molar percentage of mHtt | 47.0% | 53.2% |
| Absolute mHtt conc. | 4.3 mM | 5.1 mM |
| Molar percentage of other proteins | 53.0% | 46.8% |
| Sequestered non-mHtt protein conc. | 4.8 mM | 4.5 mM |
| **The Large Agg Magenta dot in Fig. 4a** | | |
| Molar percentage of mHtt | 56.9% | 61.5% |
| Absolute mHtt conc. | 5.6 mM | 6.4 mM |
| Molar percentage of other proteins | 43.1% | 38.5% |
| Sequestered non-mHtt protein conc. | 4.3mM | 4.0 mM |

**Table S1. Quantification of mHtt-97Q and non-mHtt proteins in aggregates.** Calculated results based on equations described above for a small aggregate (Fig. 4a, green point, 6.7 μm$^2$ and Fig. 4c, SRS (C-D)=478.1 a.u., CH/CD=11.5); a medium aggregates (Fig. 4a, red point, 19.1 μm$^2$ and Fig. 4d, SRS (C-D)=1190.3 a.u., CH/CD=5.1) and a large aggregates (Figure 4a, purple point, 36.5 μm$^2$, SRS (C-D)=1484.9 a.u, CH/CD=3.8).

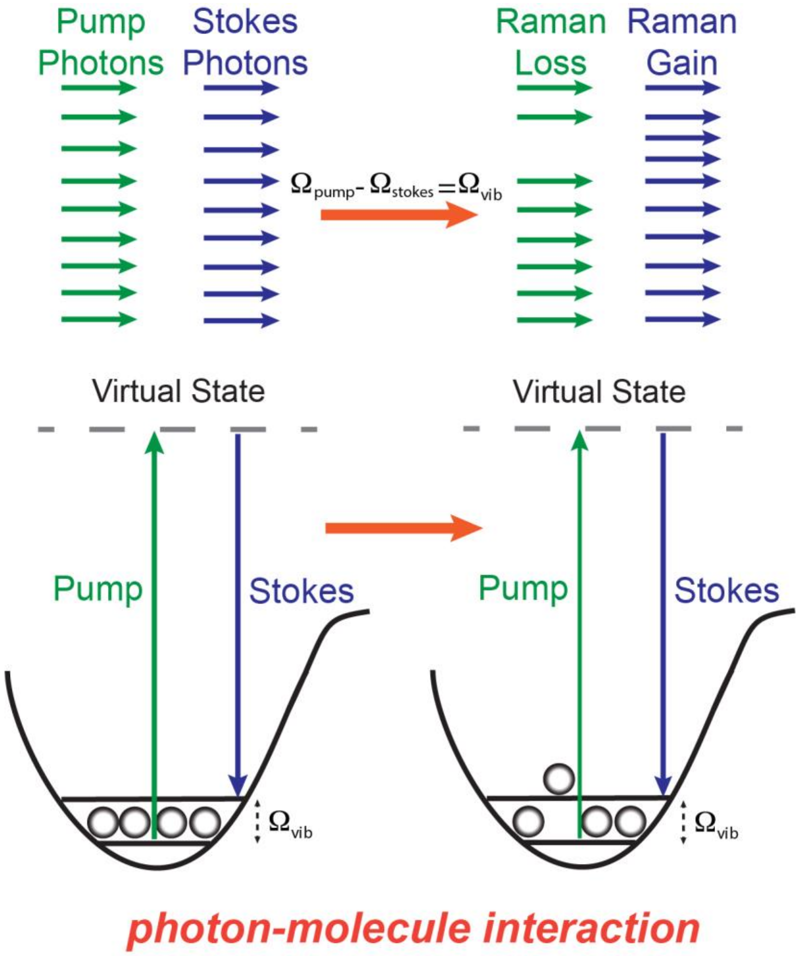

**Figure S1. Energy diagram of stimulated Raman scattering (SRS).** When the energy difference between the pump (green) and probe (blue) photons is resonant with the vibrational (Raman) transition of a C-D bond. The C-D bond would be excited to the vibrational excited state mediate by a virtual state (i.e. one grey ball, signifying C-D bond, on the left is excited to the upper level shown on the right). For each vibrational excitation of C-D bond, a pump photon is consumed (Raman Loss) and a Stokes photon is created (Raman Gain). The SRS signal is then detected as either Raman Gain or Raman loss signal. In our experiments, we adjust the laser (pump=842.8 nm, stokes=1031.2 nm) to match the vibration frequency of $\Omega_{vib}$ =2167 $cm^{-1}$ for C-D bond on Gln-$d_5$ and detect the Raman loss signal at the Pump wavelength.

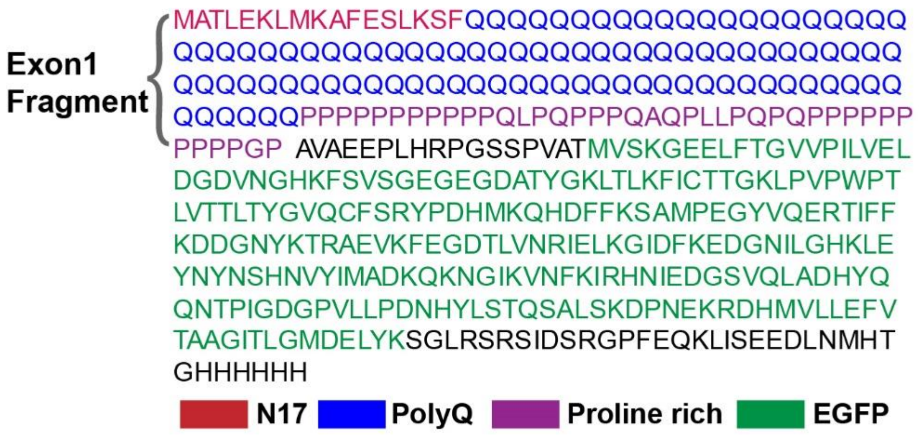

**Fig. S2. Detailed amino acid (aa) sequence information of mHtt-97Q-GFP and mHtt-97Q.** The mHtt exon1 fragment (highlighted in big parentheses), comprises of N-terminal 17 aa (red), polyQ (blue) and proline rich (purple) domain, is fused with GFP (green) by a linker region (black) followed by C-terminal myc and 6x His-tag (black).

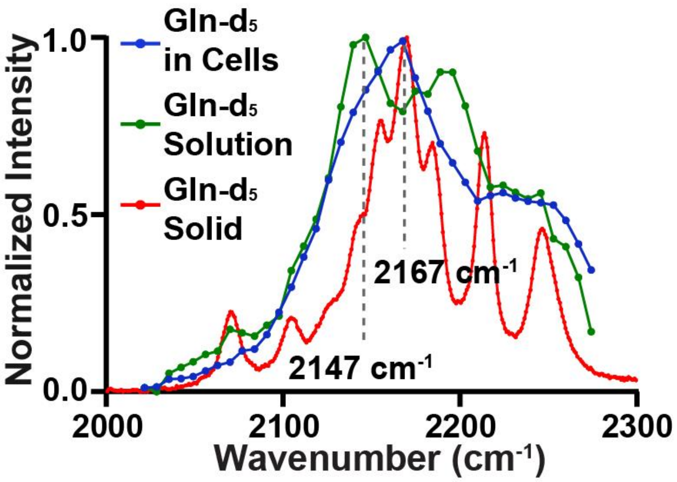

**Fig. S3. Spectral change of Gln-d$_5$ after metabolic incorporation into proteins in HeLa cells.** The Raman spectrum of Gln-d$_5$ after incorporated into cellular proteins (blue, peaked at 2167 cm$^{-1}$) presents a change in spectral line-shape compared to that of 100 mM Gln-d$_5$ solution (green, peaked at 2147 cm$^{-1}$). The shifted major peak of 2167 cm$^{-1}$ (blue) matches with the Raman peak of Gln-d$_5$ solids (red, 2167 cm$^{-1}$). Overall, the Raman spectrum of Gln-d$_5$ incorporated in cells (blue) shows combinatorial spectral features from both that of Gln-d$_5$ solutions (green) and solids (red).

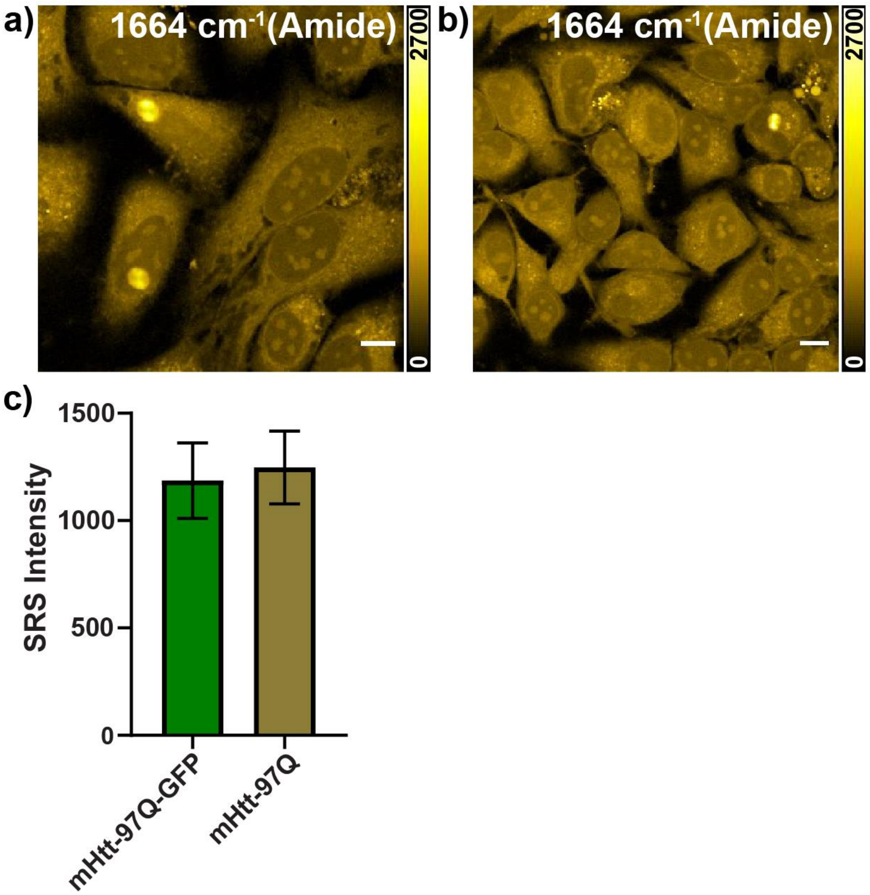

**Fig. S4. Similar average Amide I intensities for mHtt-97Q-GFP and mHtt-97Q aggregates were observed.** HeLa cells were grown in regular medium till optimal confluence and were transfected by (a) mHtt-97Q-GFP or (b) mHtt-97Q plasmids in Gln-$d_5$ medium for 24h before imaging by SRS at Amide I frequency (1664 $cm^{-1}$). (c) Quantification of average Amide I intensities for the two types of aggregates. mHtt-97Q-GFP (1186±175.8, n=10), mHtt-97Q (1248±169.4, n=14). Scale bar: 10 μm. Error bar: SD.

**Note:** Gln is a non-essential amino acid, which can be *de novo* synthesized by cells. However, the extensive demand makes it an essential supplement in culture medium. It is also reported to become a conditionally essential amino acid for cells under stress.[8,9] In both Figs. S5 and S6 below, we confirmed the Gln-$d_5$ incorporation efficiency to be near 100%, meaning that Gln from *de novo* synthesis is close to 0%.

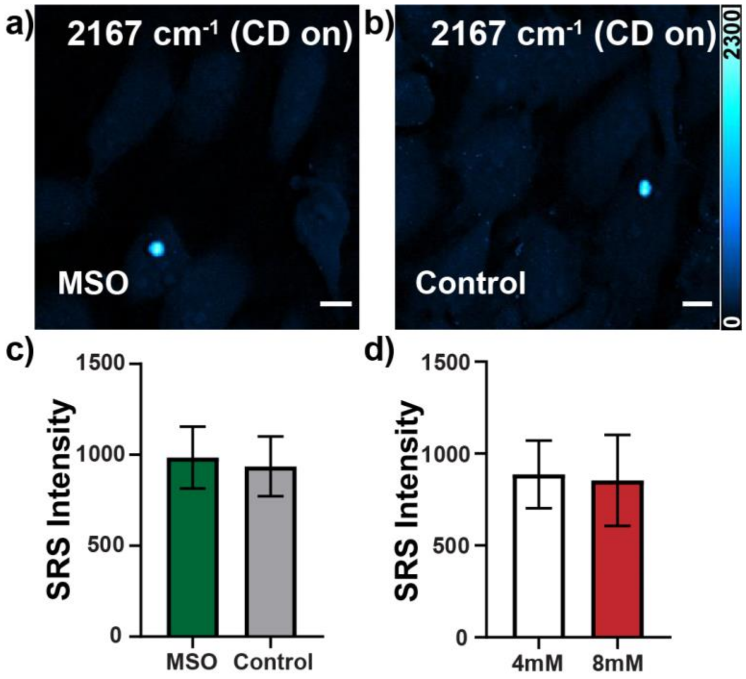

**Fig. S5. The labeling efficiency of Gln-$d_5$ for mHtt-97Q aggregates approaches 100%.** (a-b) HeLa cells transfected by mHtt-97Q was incubated with (a) or without (b) 3mM L-MSO, the Gln synthetase inhibitor[1] for inhibiting *de novo* Gln synthesis in the Gln-$d_5$ DMEM. (c) The L-MSO treated aggregates have similar intensity (984.1±160, n=9) with the non-treated ones (936.2±147, n=6). (d) Aggregates in HeLa cells, transfected by mHtt-97Q and incubated in normal DMEM (with 4mM Gln-$d_5$) or DMEM with doubled Gln-$d_5$ concentration (8mM), show no intensity difference (887.1±174, n=10 and 854.3±234, n=9 respectively). Our observations demonstrate that both inhibiting *de novo* Gln synthesis (c) and enhancing extracellular Gln concentrations (d) do not increase aggregate intensity. Our control experiments suggest that the labeling efficiency for the aggregates by Gln-$d_5$ from medium is close to 100%. Scale bar: 10 μm. Error bar: SD.

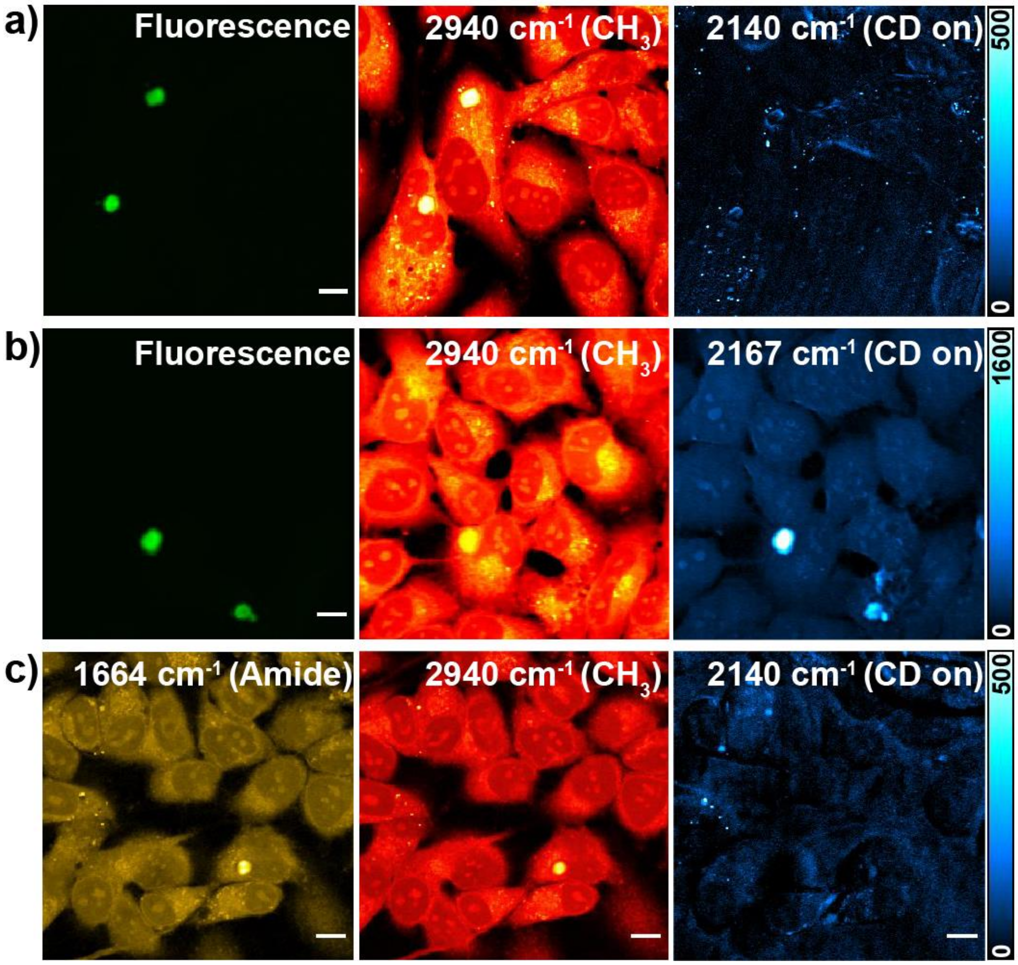

**Fig. S6. $D_7$-glucose incubation has no detectable signals in both mHtt-97Q-GFP and mHtt-97Q aggregations through *de novo* Gln synthesis.** (a-b) HeLa cells were transfected by mHtt-97Q-GFP of in (a) $d_7$-glucose medium for labeling *de novo* synthesized Gln and in (b) Gln-$d_5$ medium for Gln incorporation for 24h. $D_7$-glucose incubation shows almost no C-D signals (a, 2140 $cm^{-1}$, CD on) while Gln-$d_5$ incubation yields very bright aggregates in C-D channel (b, 2167 $cm^{-1}$, CD on). GFP is used as a guidance for aggregates due to minimum C-D signals $D_7$-glucose incubation (c) HeLa cells were first pre-incubated with $d_7$-glucose DMEM for 24h to pre-label *de novo* synthesized Gln and were next transfected with mHtt-97Q plasmid in $d_7$-glucose DMEM for another 24 h. No signals were detected for aggregates (c, 2140, CD on), suggesting a minimal level of Gln in aggregates comes from *de novo* synthesis. Scale bar: 10 μm.
